# APOE3 astrocytes can rescue lipid abnormalities and dystrophic neurites of APOE4 human neurons

**DOI:** 10.1101/2025.10.24.684364

**Authors:** Dilara O. Halim, Erika Di Biase, Amélie Rajon, Luc Jordi, Penelope J. Hallett, Ole Isacson

**Author notes:** Corresponding authors: Penelope Hallett Ole Isacson. These authors contributed equally to this work (Dilara O. Halim, Erika Di Biase).

## Abstract

Lipid abnormalities are emerging as key pathogenic mechanisms in neurodegenerative diseases such as Alzheimer’s, Parkinson’s and Lewy body dementia. Astrocytes in the brain provide APOE proteins and influence neuronal metabolism and health. Using live cell imaging and objective neurite imaging techniques, we show that following induction of cellular lipid (cholesterol and triglycerides) load by inhibiting the lysosomal cholesterol transport protein NPC1 in human neuron-astrocyte co-cultures, that human astrocytes CRISPR edited to be either APOE3 or 4 variants have different effects on rescuing dystrophic neurites, where axons and dendrites of nerve cells become disfigured. APOE3, but not APOE4 or APOEKO, astrocytes prevented cholesterol and lipid induced neurite damage in APOE4 neurons. In the media of APOE3 co-cultured astrocytes with neurons the HDL-like particles were larger and presumably more lipidated than equivalent APOE4 co-cultures. This discovery highlights that living APOE3 astrocytes control key biological mechanisms by physiologically enhancing lipid cellular homeostasis, that can rescue lipid-induced neurite structural abnormalities relevant to Alzheimer’s disease and neurodegenerative diseases.

**Significance statement:** Neurodegenerative diseases like Alzheimer’s (AD) are often defined by abnormal protein aggregates, but growing evidence points to lipid dysfunction as a key driver, especially in APOE4 carriers, the strongest genetic risk factor for AD. We developed a live cell imaging based human cell culture model using isogenic iPSC-derived neurons and astrocytes (APOE3, APOE4, or APOE knockout) to study this. By blocking cholesterol export via NPC1 inhibition, we mimicked lysosomal lipid stress and found that APOE3 astrocytes uniquely protected APOE4 neurons from forming abnormal neurite swellings. These APOE3 astrocytes produced larger HDL-like particles than APOE4 that supported neuronal lipid balance. Our results show that APOE3 astrocytes can rescue APOE4-related cellular dysfunction, offering a potential path for therapy and biomarker discovery.

## Introduction

Lipids are essential to brain function, contributing to membrane structure, energy metabolism, and regulation of signaling pathways (1, 2). Disruption of lipid homeostasis is increasingly recognized as central to the pathogenesis of neurodegenerative diseases such as Alzheimer’s disease (AD) and Parkinson’s disease (PD) (2–4). In these conditions, abnormal lipid accumulation and impaired lipid metabolism contribute to neuronal dysfunction and degeneration. Similar features occur in lysosomal storage disorders such as Niemann-Pick type C (NPC), where cholesterol trafficking defects lead to intracellular lipid accumulation and progressive neurodegeneration (5, 6).

Throughout the body, lipid-containing particles are critical for biological functions. They are classified by size and by apolipoprotein composition, which determines their specific functions and transport destinations (7). Apolipoproteins A (APOA), B (APOB), and E (APOE) interact with lipoprotein receptors to deliver and clear lipids, with most systemic clearance occurring via hepatic low-density lipoprotein (LDL) receptor-mediated pathways (8). In the brain, however, high-density lipoprotein (HDL)-like particles containing APOE are produced locally, primarily by astrocytes, and under certain conditions by microglia (9, 10). APOE is the principal lipid transporter in the central nervous system, maintaining cholesterol and lipid balance across multiple cell types (3). Humans express three APOE isoforms, APOE2, APOE3 and APOE4, that differ from each other by only single amino acid substitutions, and APOE3 being the most prevalent and APOE2 the least frequent (11). APOE4 is the strongest genetic risk factor for late-onset AD, associated with earlier dementia onset, impaired lipid metabolism, altered synaptic network function, and increased amyloid-β and tau pathology (11, 12). APOE4 has reduced cholesterol efflux capacity and produces smaller, lipid-poor APOE particles, especially in astrocytes, whose secretion of APOE depends on ABCA1-mediated lipidation (5, 13).

In grey matter and at vascular interfaces, astrocytes are central to brain lipid handling: they redistribute cholesterol and other lipids to neurons, maintain membrane integrity, and support synaptic metabolism (9). Astrocytes also provide ionic buffering, glutamate uptake, and vascular exchange through their close interactions with neurons, vessels, and other glia. These continuous, baseline homeostatic roles are vulnerable to failure under genetic or metabolic stress (4). APOE4-expressing astrocytes exhibit deficits in these functions, including reduced ABCA1-mediated lipidation and altered vesicle trafficking (5, 14). Based on such findings, we hypothesize that APOE isoform-specific differences in lipid handling alter neuron-astrocyte metabolic relationships, influencing long-term neuronal health and structure (2–4).

Across neurodegenerative disorders, disrupted cholesterol and lipid balance emerges as a recurring theme. In AD, APOE4 reduces cholesterol efflux and limits the lipidation of APOE particles (5, 15). In PD and Lewy body dementia (LBD), GBA1 mutations and age-associated loss of GBA1 function alter lysosomal glycosphingolipid metabolism and contribute to α-synuclein pathology (2, 3, 16–20). Niemann-Pick disease type C (NPC) illustrates the consequence of impaired cholesterol trafficking from lysosomes (5, 6), while in a vast number of ALS cases, altered sterol metabolism and cholesterol has been linked to motor neuron vulnerability (21). Cholesterol and lipid disturbances can modify amyloid processing of precursor protein (APP) and influence amyloid beta (Aβ) peptide balance, as reported in NPC1-inhibited and APOE-dependent models (5). Together, these observations suggest that impaired lipid and cholesterol balance converges on shared pathways of protein aggregation, mitochondrial dysfunction, and neuritic degeneration across AD, PD, LBD, NPC, and ALS.

What has been lacking scientifically are live human cell models of neuron-glia communication and their direct impact on neurites (comprised of axons and dendrites), reflecting the dystrophic changes seen in aging and likely AD (22, 23). To address this, we used isogenic human induced pluripotent stem cell (hiPSC) co-cultures of neurons and astrocytes, in which APOE4 alleles were corrected to APOE3 and vice versa. We induced intracellular cholesterol accumulation by pharmacological inhibition of NPC1, a lysosomal transporter required for cholesterol egress. This model captures how astrocytic APOE genotype influences neuronal vulnerability to lipid imbalance. APOE3 astrocytes limited cholesterol loading and reduced the appearance of abnormal neurites, while APOE4 astrocytes failed to do so. These findings highlight APOE as a key determinant of neuronal responses to cholesterol and lipid imbalances, linking astrocyte lipid handling to the emergence of neuritic pathology in AD and related disorders.

## Methods

### iPSC Lines and Maintenance

Three isogenic pairs of APOE3 and APOE4 hiPSC lines and one APOE knockout (APOEKO) line were used in this study. One isogenic set (APOE3, APOE4, and APOEKO) was purchased from Alstem, Inc. (Catalog #iPS26, iPS16, and iPS36, respectively), while the other two APOE3 and APOE4 isogenic pairs have been previously reported (15). hiPSCs were cultured on Matrigel-coated plates (Corning, #354230) in mTeSR1 medium (Stemcell Technologies, #85857) and routinely tested for mycoplasma contamination using a PCR-based detection kit (Sigma, #MP0035).

We tested the pluripotency of all lines by immunostaining for OCT4 and TRA-1-60 (Figure S1A). APOE genotype was validated by Sanger sequencing (Figure S1B,C). Genomic DNA was extracted using the DNeasy Blood & Tissue Kit (Qiagen, #69504), and the APOE region was amplified with the following primers: PCR: forward 5’-CAGGTCACCCAGGAACTGAG-3’; reverse 5’-CACCTGCTCCTTCACCTCG-3’ Sequencing: forward 5’-GCCTACAAATCGGAACTGGA-3’; reverse 5’-CTGCCCATCTCCTCCATC-3’ Sequencing was performed by Eton Bioscience (Boston, USA).

### iPSC-Derived Neuron Differentiation

iPSC-derived neurons (iNs) were generated using a previously published NGN2 overexpression protocol (24, 25), with minor modifications (Figure S2A). hiPSCs were plated at 100,000 cells/cm² on Matrigel-coated plates 1 day prior to viral transduction.

Cells were transduced with the following lentiviral constructs: pTet-O-NGN2-puro (Addgene #52047), Tet-OFUW-EGFP (Addgene #30130), and FUdeltaGW-rtTA (Addgene #19780). Transduced hiPSCs were expanded for 2 weeks, then replated at 200,000 cells/cm² in mTeSR1 with 10 μM ROCK inhibitor Y-27632 (Sigma, #Y0503) on day 0 (D0).

Media were changed sequentially: KSR medium (D1), 1:1 KSR:N2B (D2), and N2B alone (D3). On day 4 (D4), cells were dissociated with Accutase and plated at 15,000 cells/cm² in iN medium. From D1 to D10-11, doxycycline (2 μg/mL, Sigma) was added to induce NGN2 expression. Puromycin (5-10 μg/mL, Gibco) was used for selection from D2 to D10-11. B27 supplement (1:100) was added starting on D3. Cells were maintained in iN medium from D4 to D34-35 with twice-weekly half media changes.

Media compositions:

KSR medium: KnockOut DMEM, 15% KOSR, 1× MEM-NEAA, 55 μM β-mercaptoethanol, 1× GlutaMAX (Life Technologies), 100 nM LDN-193189 (Stemgent, #04-0074), 10 μM SB431542 (Tocris, #1614), 2 μM XAV (Stemgent, #04-00046).

N2B medium: DMEM/F12, 1× GlutaMAX, 1× N2 Supplement B (Stemcell Technologies), 0.3% dextrose NBM medium: Neurobasal medium, 0.5× MEM-NEAA, 1× GlutaMAX, 0.3% dextrose iN medium: NBM + B27 (1:50), supplemented with BDNF, GDNF, and CNTF (10 ng/mL each; Peprotech)

### iPSC-Derived Astrocyte Differentiation

hiPSCs were differentiated into neural progenitor cells (NPCs) using STEMdiff Neural Induction Medium with SMADi Supplement (Stemcell Technologies, #08581), following the protocol shown in Figure S2B. Cultures were maintained for 3 weeks with daily media changes and weekly passaging using Accutase. NPCs were then differentiated into astrocytes using the STEMdiff Astrocyte Differentiation Kit (Stemcell Technologies, #100-0013). Cells were cultured in differentiation medium for 3 weeks, with daily media changes during the first week and every other day thereafter. Astrocytes were then transitioned to the STEMdiff Astrocyte Maturation Medium (Stemcell Technologies, #100-0016) for 3 additional weeks, with media changed every other day and weekly passaging. Such astrocytes were used for coculture after 9 weeks of total astrocyte differentiation.

### Neuron-Astrocyte Coculture Platform and NPC1 Inhibition

For coculture experiments, iNs and hAs were generated from the same isogenic hiPSC pairs as described above. On D24-25 of iN differentiation, hAs were added to neurons at a 1:4 iN:hA ratio in 50:50 iN medium and astrocyte maturation medium. Cocultures were maintained for 1 week with twice-weekly half media changes using iN medium.

NPC1 inhibition was initiated on D31-32 by treating cells with 10 μg/mL U18666A (EMD Millipore, #662015) or PBS as vehicle control for 3 days as previously described (5) (Figures 1A, S2A). All analyses were performed following this treatment period.

**Figure 1:**
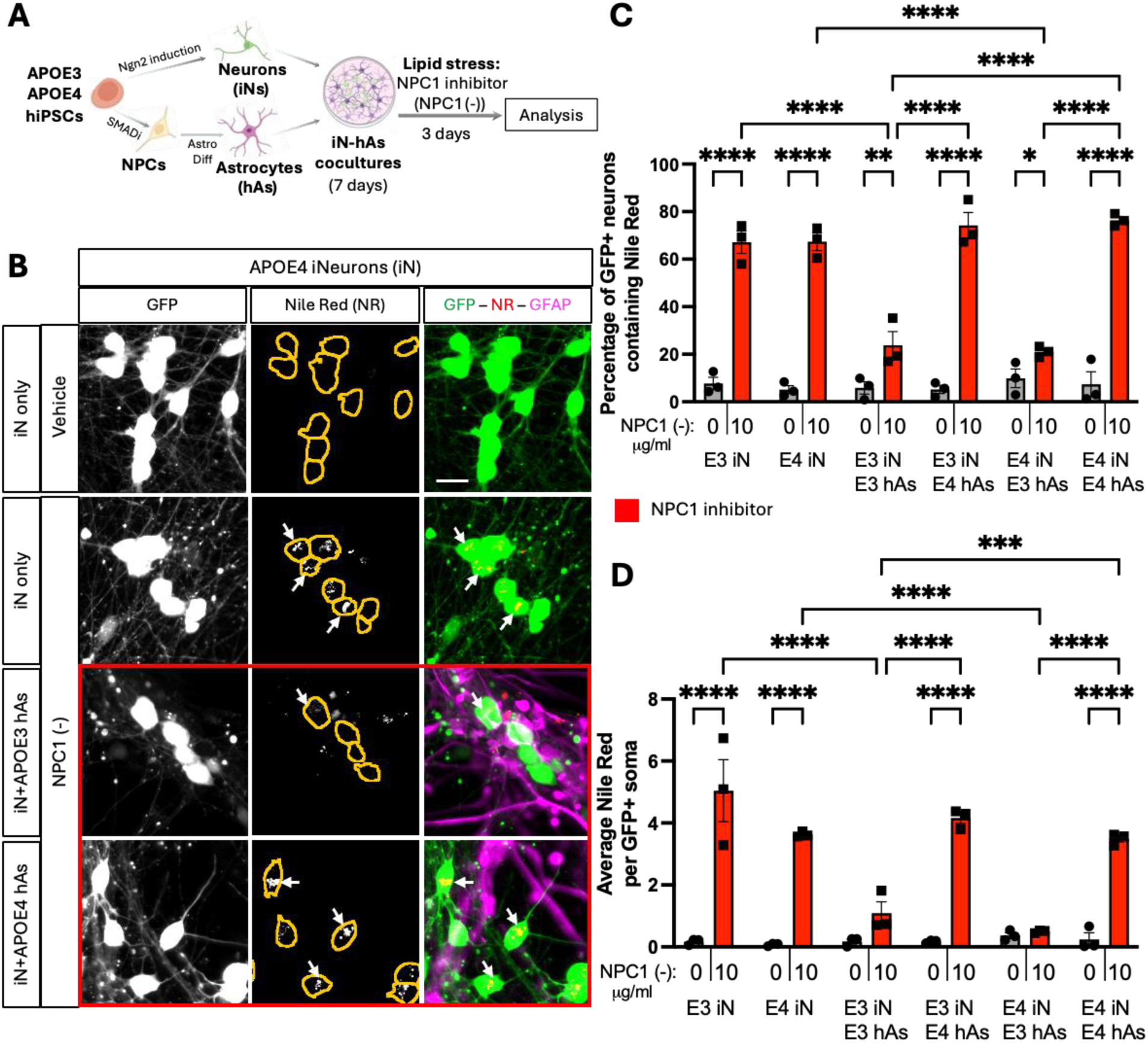
APOE3 astrocytes rescue lipid accumulation induced by NPC1 inhibition in iN. A) Schematic representation of neuron-astrocyte coculture generation (see also Figure S2) and lipid stress induction. Neurons (iN) and astrocytes (hAs) were differentiated from 3 isogenic pairs of APOE3 and APOE4 hiPSCs. After differentiation, iNs and hAs were cocultured for 1 week, followed by a 3-day treatment with either vehicle (PBS) or the NPC1 inhibitor U18666A (NPC1 (-); 10 µg/ml) to induce lipid stress. B) Representative images showing lipid accumulation in iN. Lipid droplets were visualized using Nile Red (NR) staining, with neurons expressing GFP and astrocytes labeled with GFAP immunostaining. Soma are outlined with orange lines in NR images. Scale bar: 20µm. C, D) Quantification of lipid accumulation in GFP+ iN: (C) Percentage of NR+ iN, calculated as NR iN divided by the total number of iN per image, and (D) the number of NR particles per soma per image. For each condition, at least 6 (max. 9) images were analyzed and averaged per isogenic line. Each data point represents an individual isogenic line. Statistical analysis was performed using two-way ANOVA with pairwise matching, with significant differences (p < 0.05) indicated on the graphs.

### Quantification of Neurite Abnormalities

Images were acquired using the IncuCyte® ZOOM live-cell analysis system (fluorescence dual core module 4459; green; excitation: 440-480nm; emission: 504-544nm; exported image format: TIFF) at a magnification of 20x (0.61μm/pixel). Semiquantitative analysis was performed with the ImageJ2/Fiji software package (Version 2.14.0/1.54F) (26) using the microscopy pixel classification machine learning tool Trainable Weka Segmentation (Version 4.0.0) (27). Four images were randomly selected for the analysis set-up, within which approximately 60% of the phenotypical occurrences (“swelling-abnormalities” characteristics) within each image were identified and classified (class1: phenotypical occurrence; class2: residual). These images were subsequently utilized to construct the training dataset, which was then used to train a Random Forest classifier, called Fast Random Forest (FastRandomForest-I 200-K 2-S-337290367-threads 8) (27, 28) using the following default features: Gaussian blur, Sobel filter, Hessian, Difference of Gaussian, Membrane Projections, Membrane thickness (1.0), Membrane patch size (19.0), Minimum sigma (1.0), Maximum sigma (16.0).

During phenotypical detection, small and large cellular processes, debris, or artifacts can be classified as “swelling-abnormalities phenotype”. This potential imaging noise was addressed by implementing a minimum and maximum object size parameter (min: 0.053369𝜇𝑚^2^; max: 1.174108𝜇𝑚^2^) and a minimum and maximum object circularity parameter (circularity = (4𝜋 × 𝐴𝑟𝑒𝑎)/𝑃𝑒𝑟𝑖𝑚𝑒𝑡𝑒𝑟^2^; range: 0.2-1.0) when counting phenotypical aberrations with the Analyze Particle in-house Fiji-based analytical tool (29). Thus, objects outside the specified size range were excluded from the automatic count.

The size and circularity range were determined arbitrarily based on phenotypical objects observed during classifier training and validation.

To enable rapid and consistent analysis across large image datasets, two custom Fiji macros were developed: one for batch image classification using the trained Weka model, and another for automated particle quantification. The first macro allows batch processing of multiple TIFF images using the trained Fast Random Forest classifier to generate classified probability maps. The second macro applies a fixed threshold to these output maps and executes the Analyze Particles function with the predetermined size and circularity constraints, automating the quantification of the abnormalities across batches. These macros substantially reduce analysis time and minimize user-dependent variability, ensuring reproducibility across experimental replicates.

All image analyses and quantifications of neurite abnormalities were performed in a blinded manner.

### Western blotting

Cells were harvested and lysed with cold RIPA buffer (Thermo Fisher, #PI89900) supplemented with Halt protease along with phosphatase inhibitor cocktail and EDTA (Thermo fisher, #78440). Cells were incubated on ice for 20 min after which they were sonicated (BioLogics Inc, Model 150 V) and spun down. Protein concentration of the lysates was determined using BCA assay (Thermo fisher, #23225).

20 μg protein samples were mixed with Pierce lane marker reducing sample buffer (Thermo fisher, #39000), boiled at 95 °C for 5 min, loaded onto precast 4–20% gradient Criterion Tris– HCl protein gels (Bio-Rad, #3450033) and were electrophoresed at 75 V for 10 minutes followed by 150 V for 1 h, followed by transferring onto a PVDF membrane (Bio-Rad, #1704157) at 100 V for 1 h. The membranes were incubated in blocking solution comprising 5% non-fat dry milk powder (Research Products, #M17200-500.0) in TBS-T (1 × Tris-buffered saline (Bio-Rad, #170-6435) with 0.1% Tween 20 (American Bioanalytical, #AB02038-01000)). Membranes were then incubated overnight at 4°C (on a shaker) with the following primary antibodies diluted in the blocking solution: anti-APOE (Abcam #ab183597, 1:500), and anti-beta Amyloid Precursor Protein (Thermo Fisher Scientific #CT695, 1:1000). The membranes were washed 3 times (10 min incubation on the shaker at room temperature) in TBS-T, followed by incubation in anti-rabbit IgG, HRP linked secondary antibody (Cell Signaling Technology #7074P2, 1:10000) diluted in blocking buffer, for 1 h at room temperature (on the shaker). Following another 3 washes with TBS-T (10 min incubations on the shaker at room temperature), the signals were developed using SuperSignal West Atto Ultimate Sensitivity chemiluminescent substrate (Thermo fisher, #A38556), and imaged using Chemidoc XRS with Image Lab software. Densitometry analysis was performed using Bio-Rad Image Lab software, and target protein bands were normalized to stain-free total protein bands. Stain-free total protein bands were obtained by exposing the PVDF membrane to UV light after protein transfer. For normalization, bands within the 50-37 kDa molecular weight range were selected.

### Native PAGE

Culture media were collected, centrifuged at 10,000 g for 5 min and supernatant was transferred to fresh tubes. Media samples were mixed with Native Sample Buffer (Biorad, #1610738) supplemented with G-250 Sample Additive (Invitrogen, #BN2004) and run on 4%–20% polyacrylamide tris-glycine gels in the absence of sodium dodecyl sulfate, reducing agents or sample boiling, in tris-glycine buffer supplemented with 1:20 NativePAGE Cathode Buffer Additive (Invitrogen, #BN2002), at 150 V for 15 min. The buffer was changed to tris-glycine buffer supplemented with 1:200 NativePAGE Cathode Buffer Additive and samples were continued to run at 150 V for 90 min. Native protein ladder (NativeMark Unstained Protein Standard, #LC0725, ThermoFisher) was loaded to track protein molecular weights. Proteins were transferred onto PVDF membranes as described above for western blotting. Anti-APOE (Millipore, #178479) primary antibody and anti-goat IgG, HRP (Invitrogen, #31402) secondary antibody were used to detect APOE lipoprotein bands.

The intensity of ApoE lipoprotein bands was quantified using Image Lab Software as follows: The detected ApoE lipoprotein bands were categorized into two groups based on their electrophoretic migration. Bands located between ∼700-1000 kDa were classified as large-sized lipoparticles, bands located between ∼420-700 kDa were classified as small-sized lipoparticles. To represent each lipoprotein species in a sample, the value of the specific band intensity was expressed as a percentage of the total intensity of the large and small lipoprotein bands combined. This allowed for the relative quantification of each lipoprotein species in terms of their contribution to the overall band intensity.

### ELISA

Culture media were collected as described above for Native PAGE. Media samples were diluted in respective reaction buffer 3 times for APOE, and 2 times for Aß40 and Aß42 ELISA. Extracellular APOE, Aß40 and Aß42 were detected in culture media using Apolipoprotein E Human Elisa Kit (Fisher Scientific, #EHAPOE), Human Amyloid beta (aa1-40) Quantikine ELISA kit (R&D Systems, #DAB140B), and Amyloid beta 42 Human Ultrasensitive ELISA kit (Thermo Scientific, #KHB3544) respectively following the manufacturers’ instructions.

### Immunostaining and Lipid Imaging

Cells cultured on coverslips were fixed in 4% paraformaldehyde (PFA) in PBS for 10 minutes at room temperature (RT). Following fixation, cells were washed three times with PBS and incubated for 30 minutes at RT under gentle agitation in a blocking/permeabilization solution containing 10% normal donkey serum (Jackson ImmunoResearch, #017-000-121) and 0.1% Triton X-100 in PBS (PBS-T). Cells were then incubated for 1 hour at RT with the primary antibody against GFAP (Synaptic Systems, #173004; 1:500) diluted in the same blocking/permeabilization solution, followed by three PBS washes. A 1-hour incubation at RT with Alexa Fluor 647-conjugated secondary antibody (Invitrogen; 1:500 in PBS) was performed under gentle agitation. To simultaneously stain neutral lipids, Nile Red (Thermo Fisher Scientific, #N1142; 1:1000) was included in the secondary antibody mixture. After three additional PBS washes, cells were incubated for 1 hour at RT with filipin (Sigma-Aldrich, #F9765; 0.1 mg/mL in PBS) to stain free cholesterol. Coverslips were washed again three times in PBS and mounted using ProLong™ Diamond Antifade Mountant (Thermo Fisher Scientific, #P36970).

Images were acquired at 20X magnification using a Zeiss fluorescence microscope. GFP fluorescence was used to identify neuronal soma. Cholesterol accumulation was quantified by (1) calculating the proportion of filipin-positive (filipin⁺) neurons relative to the total number of GFP-positive neurons per image, and (2) counting the number of filipin⁺ puncta per soma. Similarly, Nile Red (NR) accumulation was assessed by determining the proportion of NR⁺ neurons and the number of NR⁺ puncta per soma.

### Statistical Analysis

All statistical analyses were performed using GraphPad Prism version 10.4.2. Data are presented as mean ± standard error of the mean (SEM). Depending on the experimental design, either two-way or three-way ANOVA followed by appropriate post hoc tests was used, as specified in the figure legends. A significance threshold (α) of 0.05 was applied throughout, and *P* values less than 0.05 were considered statistically significant.

## RESULTS

### Establishment of a Neuron-Astrocyte Coculture Platform to Model Lipid Accumulation

Given the role of APOE in brain lipid metabolism and its association with neurodegeneration, we established a neuron-astrocyte coculture platform to investigate how APOE genotype impacts neuronal and glial responses to lipid accumulation. Isogenic pairs of human induced pluripotent stem cells (hiPSCs) carrying either the APOE3 or APOE4 allele (Figure S1) were differentiated into cortical glutamatergic neurons (iN) and astrocytes (hAs), and neurons labeled by GFP expression (Figure 1A, S2A, S2B).

Immunostaining confirmed efficient differentiation by lineage-specific marker expression, with iNs expressing the VGLUT2 and MAP2, and hAs expressing GFAP and S100β (Figure S2C, S2D). After differentiation, iN and hAs were maintained in coculture for 1 week prior to treatment (Figure 1A). In these cocultures, GFP+ fluorescence identified iN, while GFAP labeling confirmed hAs integration into the iN cultures (Figure 1A, S2E).

### APOE3 Astrocytes Rescue Lipid Accumulation induced by NPC1 inhibition in Neurons

Astrocytes play a key role in the regulation of neuronal lipid metabolism and lipid exchange (9). To model lipid accumulation in neuron-astrocyte cocultures and investigate how APOE genotype impacts lipid accumulation in neurons, APOE3 and APOE4 iN and iN+hAs cocultures were treated with NPC1 inhibitor U18666A (NPC1 (-); see Methods for details). NPC1 inhibition impairs lysosomal cholesterol export and induces intracellular cholesterol accumulation (5, 6). NPC1 inhibition also perturbs lipid metabolism more broadly beyond cholesterol and has been shown to trigger cellular accumulation of neutral lipids such as triglycerides and cholesteryl esters in lipid droplets (5, 30, 31). Nile Red staining, a lipophilic dye that labels both neutral lipids and phospholipids, was used to quantify neutral lipid accumulation. In iN alone, NPC1 inhibition significantly increased the percentage of Nile Red (NR)+ neurons and the number of NR+ particles per soma in both APOE3 and APOE4 iNs compared to Vehicle treatment (Veh), with no APOE genotype-dependent differences (Figure 1B-D). In iN+hAs cocultures, APOE3 hAs reduced both the proportion of Nile Red (NR)+ neurons and the number of NR+ particles per soma upon NPC1 inhibition, regardless of APOE genotype of neurons. In contrast, APOE4 hAs failed to mitigate neutral lipid accumulation (Figure 1C-D). To assess cholesterol accumulation, cells were stained with filipin, which binds to unesterified cholesterol. In iN alone, NPC1 inhibition significantly increased both the percentage of filipin+ neurons and the number of filipin+ particles per soma in APOE3 and APOE4 iNs compared to Veh., with no significant difference between APOE genotypes (Figure 2A-C). In iN+hAs, NPC1 (-) also increased the percentage of filipin+ neurons and the number of filipin+ particles per soma in both APOE3 iN+hAs and APOE4 iN+hAs compared to Veh. However, APOE4 iN+hAs exhibited significantly higher cholesterol accumulation than APOE3 iN+hAs (Figure 2A-C). Together these results highlight a genotype-specific role of APOE astrocytes in regulating neuronal lipid homeostasis.

**Figure 2.**
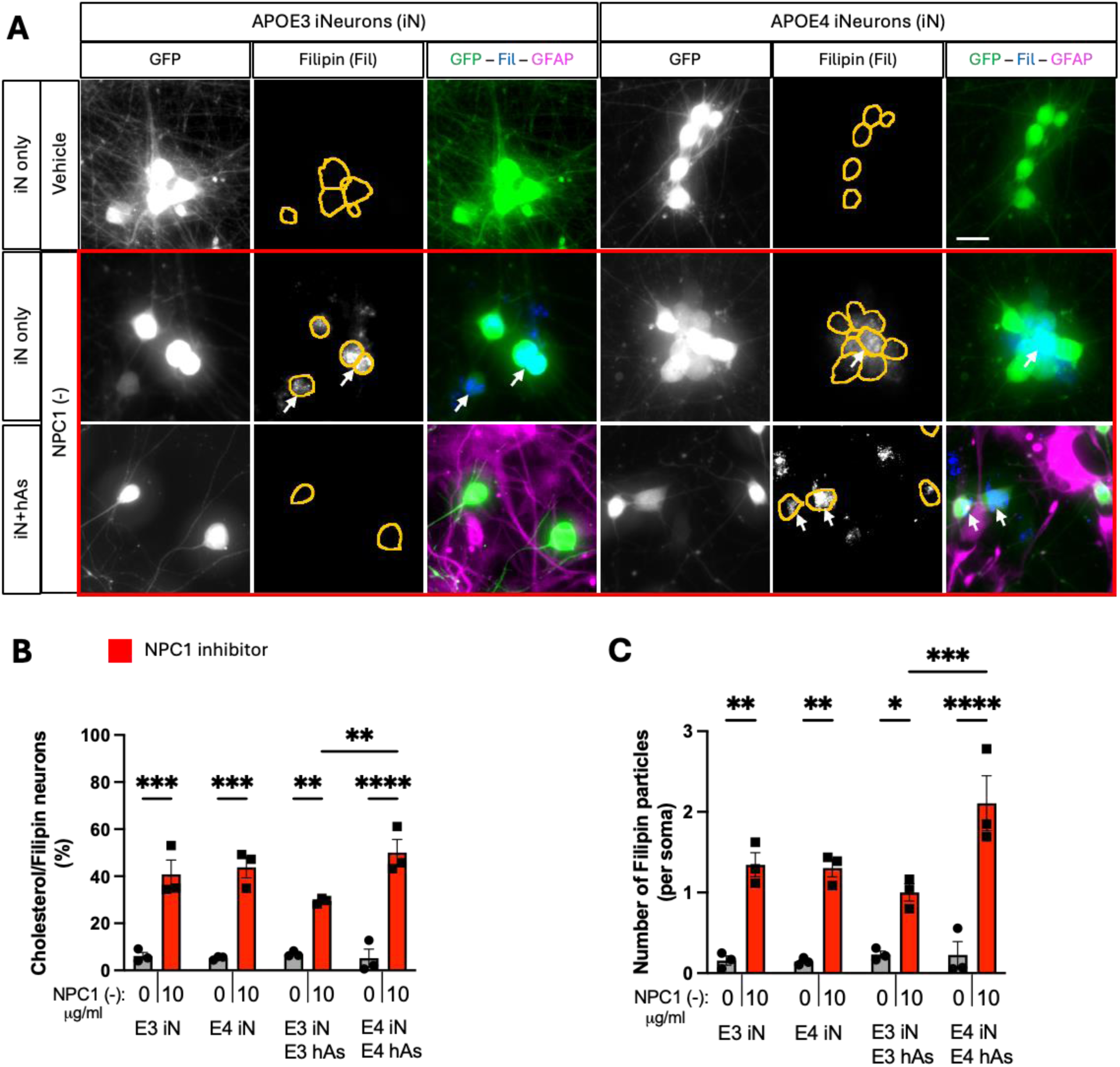
APOE3 astrocytes rescue cholesterol accumulation induced by NPC1 inhibition in iN. 3 isogenic APOE3 and APOE4 iN+hAs cocultures were maintained for 1 week and treated with either vehicle (PBS) or the NPC1 inhibitor U18666A (NPC1 (-); 10 µg/ml) for 3 days. A) Representative images showing cholesterol accumulation in iN. Cholesterol accumulation was visualized using Filipin staining, with neurons expressing GFP and astrocytes labeled with GFAP immunostaining. Soma are outlined with orange lines in Filipin images. Scale bar: 20 µm. B, C) Quantification of cholesterol accumulation in GFP+ iN: (B) Percentage of Filipin+ iN, calculated as Filipin+ iN divided by the total number of iN per image, and (C) the number of Filipin+ particles per soma per image. For each condition, 9 images were analyzed and averaged per isogenic line. Each data point represents an individual isogenic line in panels (B and C). Statistical analysis was performed using two-way ANOVA with pairwise matching, with significant differences (p < 0.05) indicated on the graphs.

### APOE3 Astrocytes Rescue Neurite Abnormalities Induced by NPC1 Inhibition

We examined neurite morphology following NPC1 inhibition. Phase-contrast microscopy revealed neurite abnormalities upon induction of lipid challenge, characterized by discrete swellings along neurites, which corresponded with GFP+ puncta (Figure 3A), allowing GFP fluorescence to serve as a quantifiable readout of neurite abnormalities. Additional staining with phalloidin showed that while F-actin was uniformly distributed along neurites at baseline, NPC1 inhibition led to F-actin accumulation in neurites, partially colocalizing with GFP+ puncta, further defining the nature of these neurite abnormalities (Figure 3B).

**Figure 3:**
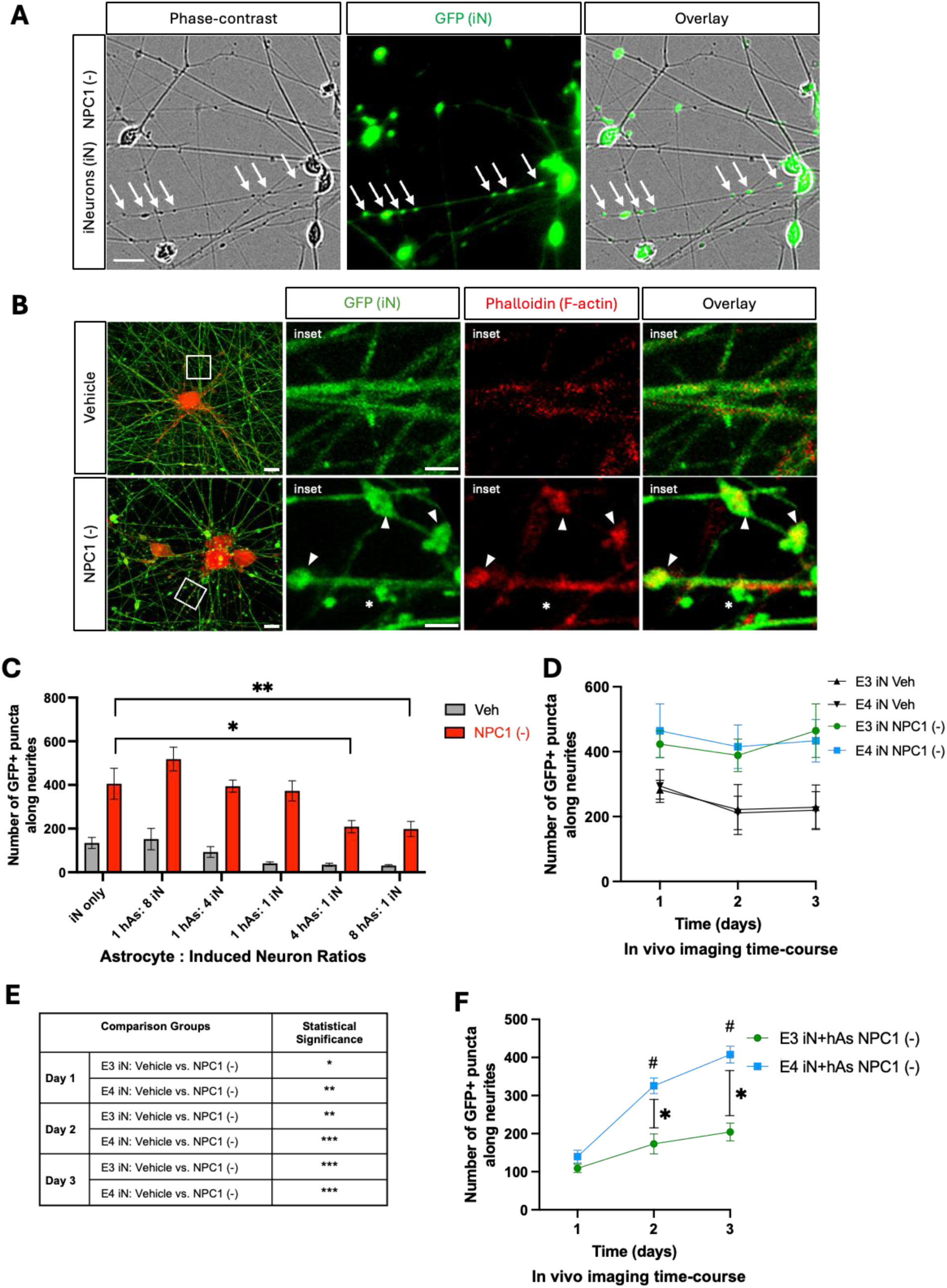
NPC1 inhibition induces neurite abnormalities in iN and iN+hAs. **Neurons (iN)** and astrocytes (hAs) were differentiated from a human iPSC line and cocultured for 1 week following differentiation. Cultures were then treated for 3 days with either vehicle (PBS) or the NPC1 inhibitor U18666A (NPC1 (-); 10 µg/ml) to induce lipid stress. (A) Representative images of iN captured using phase-contrast microscopy, GFP fluorescence, and their overlay. Neurite abnormalities, indicated by white arrows, seen as swellings observed in the phase-contrast image correspond with GFP+ puncta, which were therefore quantified as a proxy for neurite abnormalities. Scale bar: 20 µm. (B) Representative images of iN labeled with GFP (green) and phalloidin staining for F-actin (red), with the merged overlay shown. Rectangular insets indicate regions displayed at higher magnification on the right. Arrowheads highlight neurite abnormalities colocalizing with phalloidin, while asterisks mark neurite abnormalities without colocalization. Scale bar: 2 µm. (C) iN were cultured alone or cocultured with hAs at varying hAs: iN ratios, and neurite abnormalities were quantified as GFP+ puncta along neurites. For each condition, at least 15 images were analyzed. (D-E) Neurite abnormalities, quantified as GFP+ puncta along neurites, were analyzed in iN (D, E). (F) iN+hAs cocultures from 3 isogenic pairs of APOE3 and APOE4 lines following NPC1 inhibition. Neurite abnormalities were quantified at 1-, 2-, and 3-days post-treatment. Data for each condition were averaged from 3 independent experiments per isogenic line. For each condition, at least 15 images were analyzed per independent experiment (full differentiation of all isogenic lines) and data for each condition were averaged from 3 independent experiments per isogenic line. Each data point represents average of 3 isogenic APOE3 or APOE4 line. Statistical analysis was performed using two-way ANOVA for panels (C, D, and F), and three-way ANOVA for panel (E), with significant differences (p < 0.05) indicated on the graph. For panel (C), treatment with NPC1 inhibitor significantly increased neurite abnormalities compared to vehicle in all conditions (p < 0.0001 for each), though these differences are not marked on the graphs. For panel (D), significant differences are shown in panel (E). For panel (F), asterisks indicate significant differences between APOE3 and APOE4; hash symbols indicate significant changes relative to day 1.

Prior to investigating APOE genotype effects, the neuron-astrocyte coculture conditions were optimized. Neurons were plated with increasing numbers of astrocytes and treated with NPC1 inhibitor (see Methods for details). While NPC1 inhibition significantly increased the number of GFP+ puncta in all conditions, cocultures with astrocytes at 4 hAs:1 iN and 8 hAs:1 iN ratio significantly reduced neurite abnormalities compared to iN monocultures (Figure 3C). Since the 4 hAs:1 iN ratio is consistent with physiological astrocyte:neuron ratios observed in the human brain (32), this condition was selected for subsequent experiments. Notably, coculture of neurons with human fibroblasts at the same ratio (4 fibroblasts:1 iN) failed to rescue neurite abnormalities induced by NPC1 (data not shown), confirming the specificity of astrocyte-mediated neuroprotection.

A time-course analysis of neurite abnormalities revealed when these abnormalities first emerged and when differences between APOE3 and APOE4 cultures began to appear (see Methods for details). NPC1 inhibition significantly increased neurite abnormalities in both APOE3 and APOE4 iN compared to Veh, starting on day 1 of the NPC1 inhibition treatment and continuing over the 3-day period. No significant differences were observed between APOE3 and APOE4 groups (Figure 3D,E). In contrast, APOE3 and APOE4 iN+hAs cocultures showed comparable levels of neurite abnormalities at day 1, but their trajectories diverged thereafter (Figure 3F). While APOE3 iN+hAs did not show a significant increase in neurite abnormalities across the treatment period, APOE4 iN+hAs exhibited a robust and progressive increase, reaching significance by days 2 and 3. These findings indicate that the protective effects of APOE3 astrocytes were sustained over time, whereas APOE4 astrocytes failed to prevent cumulative lipid accumulation (Figure 3F).

To investigate the impact of astrocytic APOE genotype on NPC1 inhibitor-induced neurite abnormalities in neuron-astrocyte cocultures, APOE3 and APOE4 iN were cocultured with either APOE3 or APOE4 hAs (Figure 4A). At the end of 3-day treatment period, APOE3 and APOE4 iN exhibited similar numbers of GFP+ puncta at baseline (Veh). Coculturing iNs with hAs reduced the neurite abnormalities in both APOE3 and APOE4 cultures at baseline (Figure 4B). Following NPC1 inhibition, the number of GFP+ puncta significantly increased across all conditions, showing that NPC1 inhibitor-induced lipid stress exacerbates neurite abnormalities. However, regardless of neuronal APOE genotype, APOE3 hAs significantly reduced the number of GFP+ puncta induced by lipid stress, whereas APOE4 hAs did not (Figure 3B).

**Figure 4:**
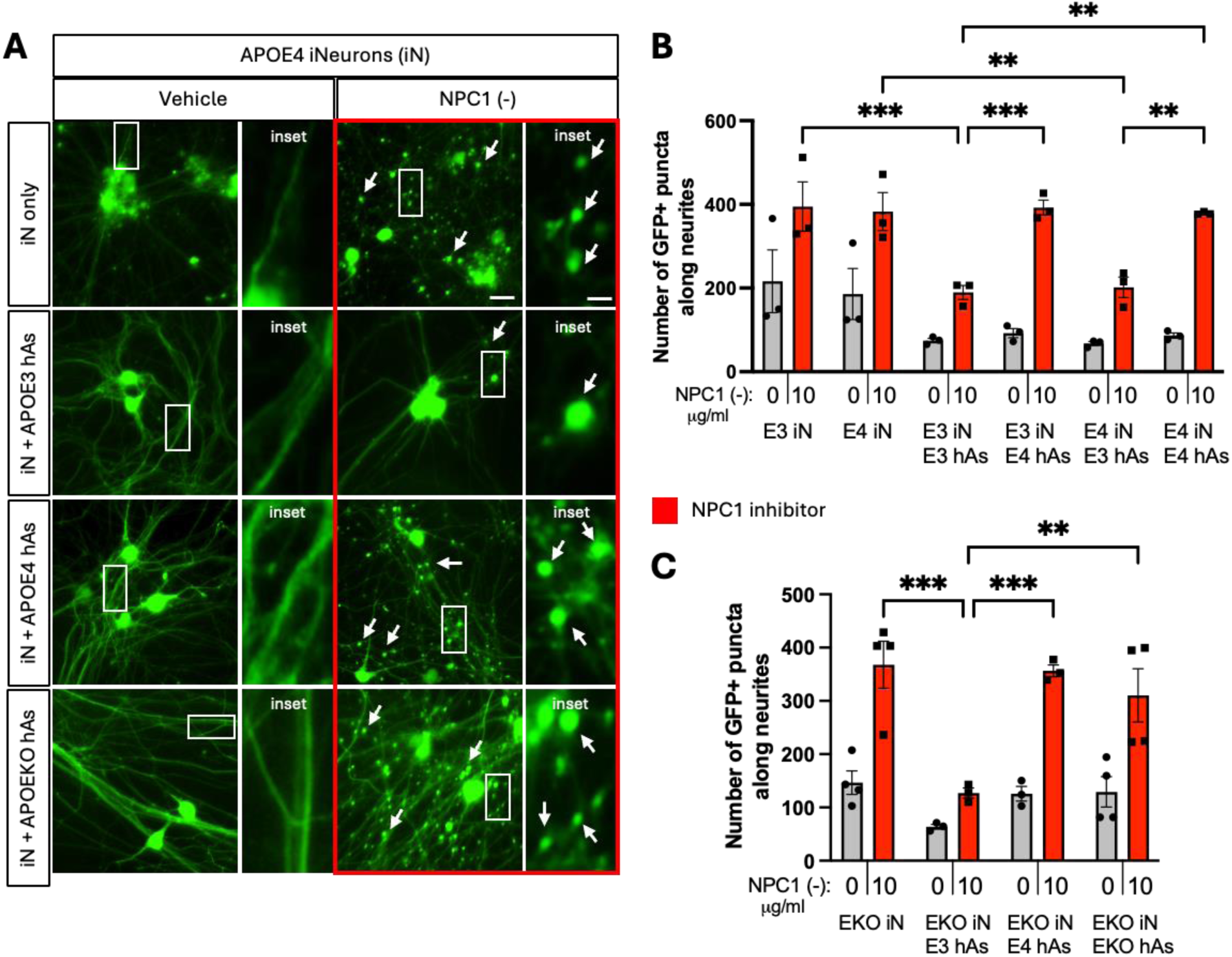
C**o-culture of APOE4 neurons with APOE3 astrocytes reduced neurite abnormalities induced by NPC1 inhibition.** 3 isogenic APOE3 and APOE4 iN+hAs cocultures were maintained for 1 week and treated with either vehicle (PBS) or the NPC1 inhibitor U18666A (NPC1 (-); 10 µg/ml) for 3 days. (A) Representative images of neurite abnormalities, indicated by white arrows in APOE3 and APOE4 iN and iN+hAs cultures treated with vehicle or NPC1 (-). Rectangular insets indicate regions displayed at higher magnification on the right. Scale bar: 20 µm, scale bar in higher magnification images: 5 µm. (B) Quantification of neurite abnormalities, measured as number of GFP+ puncta along neurites per image, in APOE3 and APOE4 iN and iN+hAs cultures treated with vehicle or NPC1 (-). (C) Quantification of neurite abnormalities in APOEKO iN and iN+hAs cultures under the same conditions. For each condition, at least 15 (max. 25) images were analyzed per independent experiment (full differentiation of all isogenic lines) and data for each condition were averaged from at least 3 independent experiments per isogenic line. Each data point represents an isogenic line in panel (C) and an independent experiment in panel (D). Statistical analysis was performed using two-way ANOVA with pairwise matching, with significant differences (p < 0.05) indicated on the graph. NPC1 (-) treatment significantly increased neurite abnormalities compared to vehicle in all conditions, though these differences (p<0.0001 for each condition) are not shown on the graph.

To determine how essential APOE was for these observed neurite abnormalities, APOE knockout (KO) iN and iN+hAs were also analyzed alongside their isogenic APOE3 and APOE4 counterparts. As seen with APOE4 hAs, APOE KO hAs failed to reduce the number of GFP+ puncta under lipid and cholesterol accumulation (Figure 4C).

### APOE3 Astrocytes Have Higher Levels of APOE and Higher Levels of Large APOE Particles at Baseline and upon NPC1 Inhibition

Using native PAGE followed by APOE western blotting, the levels of small and large extracellular APOE particles were determined. Levels of small extracellular APOE particles remained unchanged across all conditions and genotypes, both at baseline and following NPC1 inhibition. In contrast, differences emerged in the large extracellular APOE particles. At baseline (Figure 5A,B), cocultures with APOE3 astrocytes exhibited significantly higher levels of large extracellular APOE particles compared to those with APOE4 astrocytes. This difference became more pronounced under lipid challenge, as APOE3 astrocyte cocultures significantly increased large extracellular particle levels in response to NPC1 inhibition, while APOE4 cocultures did not show a similar response (Figure 5A,B).

**Figure 5:**
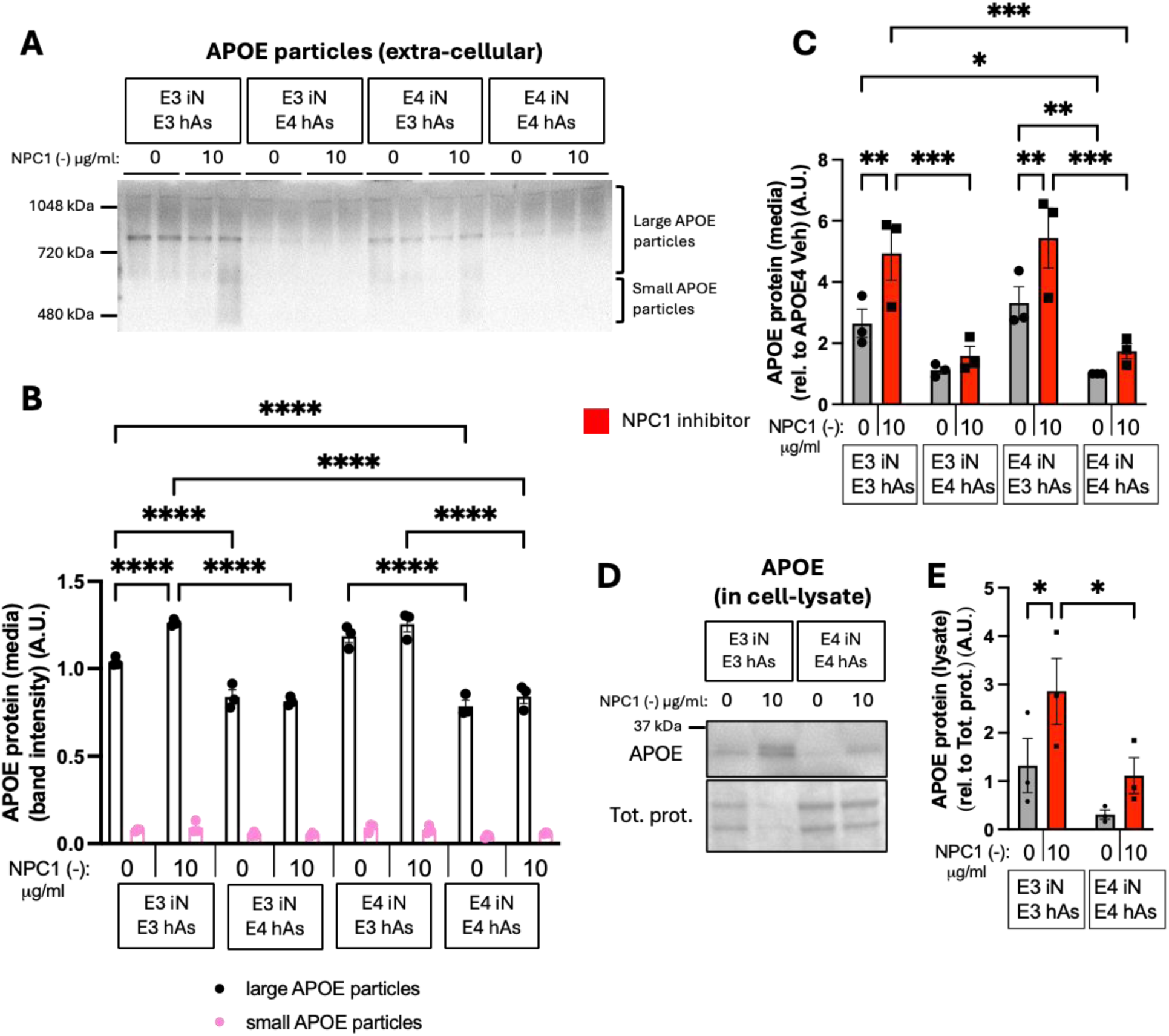
A**P**OE3 **astrocytes have higher levels of APOE and higher levels of large APOE particles at baseline and upon NPC1 inhibition.** 3 isogenic APOE3 and APOE4 iN+hAs cocultures were maintained for 1 week and treated with either vehicle (PBS) or the NPC1 inhibitor U18666A (NPC1 (-); 10 µg/ml) for 3 days. (A) Representative native PAGE followed by APOE western blot showing large and small APOE particles in the culture media. (B) Quantification of large and small APOE particle band intensity from native PAGE blots. (C) Total APOE levels measured in culture media using an APOE ELISA assay. (D) Representative western blot showing total APOE levels in iN+hAs lysates. (E) Quantification of APOE band intensity from western blots. APOE levels were normalized to total protein levels. Data for each condition were averaged from at least 3 independent experiment (full differentiation of all isogenic lines) per isogenic line in panels (B and C), and from 2 independent experiment per isogenic line in panel (E). Each data point represents an isogenic line in panels (B, C, and E). Statistical analysis was performed using two-way ANOVA with pairwise matching. Significant differences (p < 0.05) are indicated on the graphs.

It is important to examine both total extracellular and intracellular APOE levels. At baseline, APOE3 and APOE4 iN cocultured with APOE3 hAs exhibited significantly higher extracellular APOE levels compared to those with APOE4 hAs cocultures. Upon NPC1 inhibition, cocultures with APOE3 hAs further increased extracellular APOE levels, whereas cocultures with APOE4 hAs showed no change (Figure 5C). Moreover, NPC1 inhibition significantly increased intracellular APOE levels in APOE3 iN+hAs, but not in APOE4 iN+hAs, leading to significantly higher intracellular APOE levels in APOE3 iN+hAs compared to APOE4 iN+hAs following NPC1 inhibition (Figure 5D,E). These findings demonstrate that APOE3 hAs exhibited higher levels of APOE and higher levels of large APOE particles at baseline and in response to lipid stress, which may contribute to their neuroprotective function and role in maintaining neuronal lipid homeostasis.

### NPC1 Inhibition Alters Levels of APP and its Cleavage Products in APOE3 and APOE4 Neurons and Neuron plus Astrocyte Co-Cultures

Given the genetic association between APOE risk and amyloidogenic proteins such as amyloid precursor protein (APP), we next examined whether NPC1 inhibition also impacted APP metabolism. To evaluate the impact of NPC1 inhibition and lipid stress on Aβ processing, extracellular Aβ40 and Aβ42 levels were measured in iN and iN+hAs, and the Aβ42/40 ratio was calculated. In iN+hAs, the Aβ42/40 ratio significantly increased following NPC1 inhibition in both APOE3 and APOE4 cocultures (Figure 6A) primarily driven by a reduction in Aβ40 levels. In APOE3 and APOE4 iN, the Aβ42/40 ratio did not change upon NPC1 inhibition (Figure 6F). Under conditions of lipid accumulation, extracellular Aβ40 levels significantly decreased in both APOE3 and APOE4 iN (Figure 6B) and in iN+hAs (Figure 6C). Extracellular Aβ42 levels showed no change across most conditions except for a significant decrease in APOE4 iN (Figure 6D), and a significant increase in APOE3 iN+hAs (Figure 6E) upon NPC1 inhibition. We also determined the full-length APP (APP-FL) and its cleavage products APP C-terminal fragments (APP-CTF). APP-FL and APP-CTF levels significantly increased in APOE4 iN following NPC1 inhibition, but showed no significant change in APOE3 iN (Figure 6G-I). In iN+hAs, APP-FL and APP-CTF levels were increased in response to NPC1 inhibition in APOE3 and APOE4 cocultures, leading to significantly higher levels of both proteins in APOE3 iN+hAs compared to APOE4 iN+hAs (Figure 6J-L).

**Figure 6.**
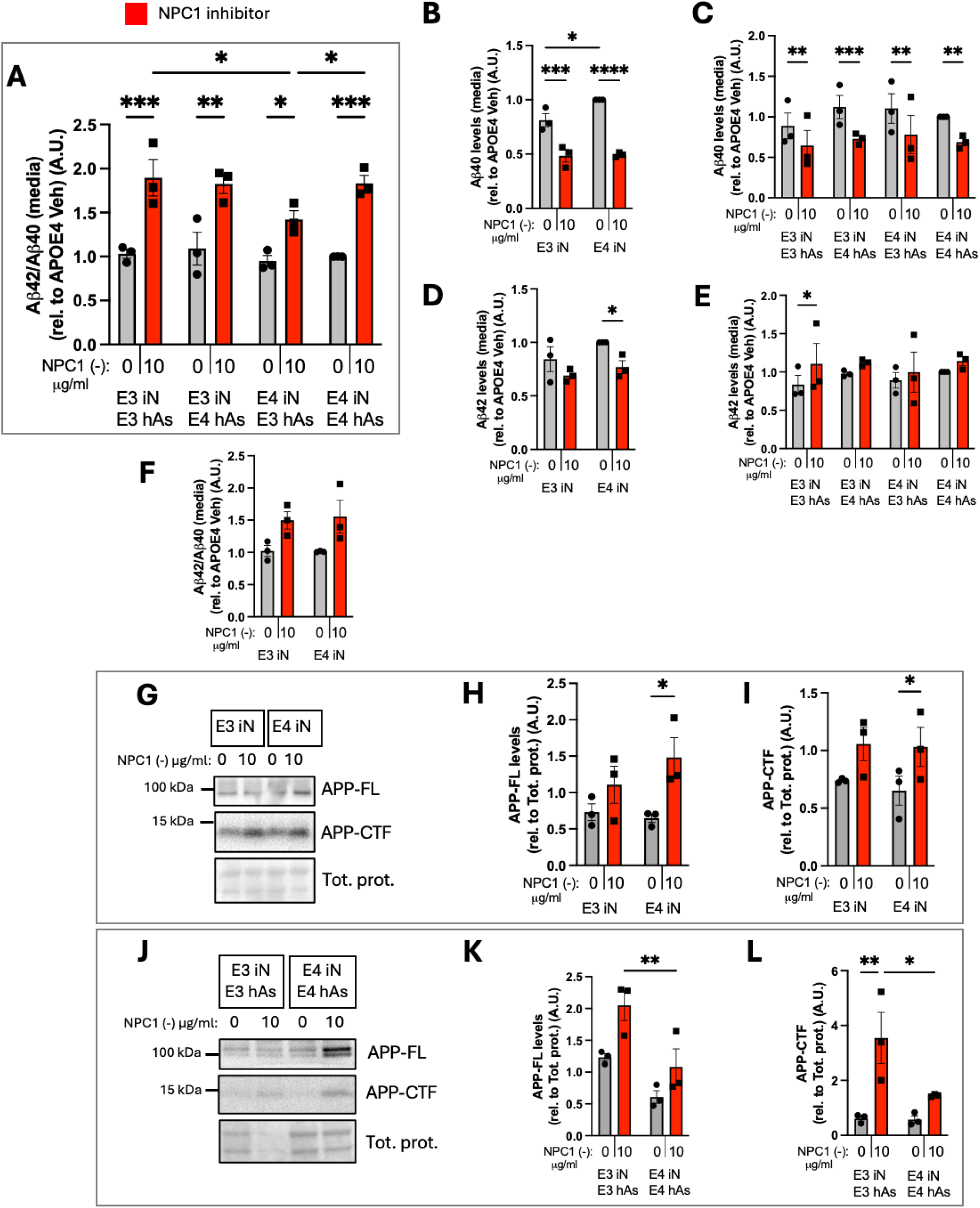
NPC1 inhibition alters levels of APP and associated peptides in APOE3 and APOE4 iN and iN-hAs. 3 isogenic APOE3 and APOE4 iN+hAs cocultures or iN were maintained for 1 week and treated with either vehicle (PBS) or the NPC1-inhibitor U18666A (NPC1 (-); 10 µg/ml) for 3 days. (A-F) Total extracellular Aβ42 and Aβ40 levels were measured in culture media using ELISA assays. Aβ42/40 ratio is shown in (A) iN+hAs and (F) iN. Aβ40 levels are shown in (B) iN and (C) iN+hAs, Aβ42 levels are shown in (D) iN and (E) iN+hAs. (G, J) Representative western blots showing full-length APP (APP-FL) and C-terminal APP fragment (APP-CTF) in (G) iN and (J) iN+hAs. (H-I, K-L) Quantification of APP-FL and APP-CTF band intensity from western blots is shown in (H-I) iN and (K-L) iN+hAs. Protein levels were normalized to total protein. Data represent an average of at least 3 independent experiments per line in panels (A-F) and average of 2 independent experiments per line in panels (H-I and K-L). Each data point represents an isogenic line. Statistical analysis was performed using two-way ANOVA with pairwise matching. Significant differences (p < 0.05) are indicated on the graphs.

Together, these findings indicate that NPC1 inhibition increased APP levels and altered Aβ processing in both iN and iN+hAs.

## DISCUSSION

Using isogenic human iPSC-derived neurons and astrocytes carrying either the APOE3 or APOE4 allele, a coculture system was created to investigate how APOE genotype impacted neuronal and glial responses to lipid accumulation. APOE3 astrocytes significantly reduced neuronal cholesterol and lipid accumulation under NPC1 inhibition, whereas APOE4 astrocytes failed to do so. Quantitative imaging-based assay demonstrated the time course of morphological, biochemical and structural changes in human APOE3 and APOE4 neurons co-cultured with their isogenic counterparts or CRISPR-edited counterpart astrocyte version (APOE3 with 4 and vice versa). This system enabled a key discovery -- in response to lipid and cholesterol accumulation under NPC1 inhibition, neurite abnormalities, including focal swellings and associated cytoskeletal changes, could be rescued by APOE3 astrocytes, but not APOE4. These neuritic changes in vitro are reminiscent of pathological dystrophic neurites and axonal swellings described in postmortem brains of patients with neurodegenerative diseases, such as AD (22, 23, 33).

In these experiments, neurite abnormalities were among the earliest detectable signs of neuronal phenotypic changes following NPC1 inhibition. Notably, the neuronal abnormalities were dependent on the astrocyte APOE genotype, emphasizing cell to cell interactive mechanisms. These results establish that astrocyte APOE status, rather than neuronal APOE, modulates neuronal responses to lipid load. The finding that APOE3 astrocytes normalize neuritic abnormalities present in APOE4 neurons highlights the critical biological significance of neuron and glia interactions in the context of lipid dysregulation.

### Human Astrocytic APOE Genotype Determines Neuronal Lipid Handling and Cell Structural Vulnerability

The current findings demonstrated that neuronal abnormalities, including neurite swellings, were dependent on the astrocyte APOE genotype. This effect was observed regardless of the neuronal APOE genotype, highlighting the astrocytic control of neuronal lipid handling. These observations are consistent with the view that astrocytic mechanisms can determine the outcomes of neuronal cell integrity and relative vulnerability to degenerative processes (14).

Astrocytic HDL-like particles are central to maintaining brain lipid balance, analogous to the liver’s role in peripheral lipid metabolism (7, 8, 34). For context, in the blood circulation, APOE-associated HDL particles exchange cholesteryl esters with larger, cholesterol-and triglyceride-rich LDL particles via cholesteryl ester transfer protein (CETP) (35). In the brain, lipid exchange occurs primarily through APOE-containing HDL-like particles and ABCA1 transporters. Both excess and deficiency in HDL-like particle activity can disrupt synaptic maintenance and membrane repair (11, 36). APOE4 disrupts this equilibrium by producing poorly lipidated particles and altering their composition, thereby reducing cholesterol redistribution. In support of this broader concept, suppression of γ-secretase in human neurons reduces cholesterol levels and impairs synaptic transmission (37). APOE4 is also the strongest genetic risk factor for cerebral angiopathy (CA), a condition in which poorly lipidated particles and associated cholesterol, together with Aβ-lipoprotein complexes, can become trapped within vessel walls, altering vascular integrity and triggering a local inflammatory response. Notably, APOE4 astrocytes secrete higher levels of inflammatory cytokines such as IL-6 and TNFα upon activation, further amplifying the pro-inflammatory milieu (38, 39). Our previous studies (17, 40–42) have shown that glial activation and inflammatory cytokine networks, whether initiated by bacterial-or viral-like stimuli or lysosomal lipid accumulation, can prime neuronal circuits for degeneration, lowering the threshold for synaptic and structural failure. The convergence of impaired lipid homeostasis with vessel wall-driven inflammation may therefore represent a key mechanism by which APOE4 increases neurodegenerative risk.

### Biological Mechanisms Involved in Astrocyte-Neuron Lipid Exchange Producing Neuritic and Other Brain Degenerative Changes

Our findings reveal that APOE genotype critically modulates neuritic vulnerability under lipid stress. APOE4 astrocytes, like APOE knockout astrocytes, failed to rescue lipid-induced neurite abnormalities, indicating a loss-of-function relative to APOE3. These abnormalities likely reflect cholesterol’s central role in maintaining membrane and cytoskeletal organization. NPC1 inhibition induced discrete F-actin-rich swellings along neurites, visualized as GFP+ puncta, representing structural abnormalities linked to impaired cytoskeletal regulation. Our finding that neuritic swellings contain F-actin parallels prior observations that lipid-binding proteins (e.g., α-synuclein) form actin-positive dystrophic structures (43, 44).While APOE3 and APOE4 neurons alone were equally affected following NPC1 inhibition, co-culture with APOE3 astrocytes, but not APOE4 astrocytes (or APOEKO astrocytes), markedly rescued these abnormalities, clearly demonstrating that APOE3 expression in astrocytes is sufficient to reverse these neuritic phenotypes. These results unequivocally show that astrocytic APOE is essential for neuronal support under conditions of lipid imbalance. Although the F-actin-positive swellings we observe are not identical to the dystrophic neurites present in human AD, the data suggest that similar upstream factors, including defective cholesterol handling, altered APP cleavage, and isoform-specific APOE effects, can converge on neuritic vulnerability. This model may therefore represent an early stage of neuronal injury relevant to the formation of dystrophic neurites in vivo. Consistent with this idea, neuropathological studies in NPC brain have shown that tau-positive neurofibrillary tangles develop in the absence of amyloid plaques, reinforcing the view that lipid dysregulation alone is sufficient to trigger cytoskeletal pathology in vulnerable neurons (45). Incorporating additional markers of neuritic pathology in future studies will help to strengthen this connection.

Interestingly, the APOE3 astrocyte as a living mechanism provided the support and prevention of the dystrophic neurites. Previously, we and others have demonstrated that cellular presentation of a trophic protein can be neuroprotective in neurodegenerative in vivo modeling, which is comparably more effective than protein infusion methods (46). Future studies can address how astrocytic presentation of APOE3 is similar or different to other methods of APOE3 delivery in vitro or in the living brain.

Prior work has shown that APOE4 astrocytes display impaired ABCA1-mediated cholesterol efflux, reduced lipid droplet dynamics, and smaller, poorly lipidated particles (47, 48). Di Biase et al. (5) further established that APOE4 astrocytes produce smaller extracellular APOE complexes with lower cholesterol content compared to APOE3, providing direct evidence that APOE4-associated particle formation defects persist even under basal conditions. These molecular deficits are significant, given that APOE-lipid complexes serve as critical vehicles for intercellular lipid transport in the brain (49).

Cholesterol transport, location and sensing is pivotal for cell biological functions. The current findings emphasize the pivotal role of astrocytic APOE, not only in regulating glial lipid handling, but also in shaping neuronal responses to lysosomal lipid and cholesterol stress. While neurons cultured in isolation can upregulate APOE and activate limited cholesterol biosynthesis (50, 51), these responses are compensatory and insufficient to sustain their extensive membrane and neurite networks. This limitation reflects an evolutionary shift toward neuronal reliance on astrocytic delivery of cholesterol-rich, APOE-containing lipoprotein particles. Consistent with this, APOE4 astrocytes impair cholesterol efflux and promote intracellular lipid accumulation (15, 47, 52). The neuron-astrocyte co-culture system used in the current study allowed direct observation of how these astrocytic genotype differences influence lipid-laden neurons. Lipid buildup in this context localizes to lysosomes but also extends to other intracellular compartments, suggesting that defects in trafficking and cholesterol sensing are not confined to a single organelle but reflect a more systemic disruption of lipid handling (53).

These observations align with prior work showing that NPC1 inhibition triggers a maladaptive feedback loop in cholesterol sensing, where reduced cholesterol trafficking to the ER upregulates HMGCR and suppresses ABCA1, exacerbating intracellular lipid accumulation (5). In APOE4 astrocytes, this feed-forward dysregulation may be compounded by intrinsic lipid export deficits, resulting in failure to restore homeostasis.

Mechanistically, APOE4 misfolding impairs ABCA1-mediated lipidation, with the protein retained in the ER and targeted for degradation (54). APOE4 also disrupts endosomal recycling, reducing secretion and compromising membrane repair and lipid redistribution (47, 53, 54). These trafficking defects converge with impaired extracellular APOE particle formation to produce a multifactorial loss of function in APOE4-expressing astrocytes.

Beyond lipid export, APOE4 astrocytes display broader impairments that may compromise their neuroprotective capacity. These include reduced neurotrophic support, diminished glutamate uptake, and altered mitochondrial structure (38, 55, 56), together with changes in lipid-regulatory and synaptic support gene expression (15). Conditioned media experiments further show that APOE4 astrocytes increase APP expression and Aβ42 secretion in APOE3 neurons, likely through cholesterol-driven lipid raft expansion (57), while failing to promote synaptic marker expression (PSD-95, SNAP-25) or enhance neuronal viability to the extent achieved by APOE3 astrocytes (14). In contrast, APOE3 astrocyte-conditioned media reduces neuronal lipid accumulation and provides trophic support, even to APOE4 neurons (14).

APOE3 astrocytes generate higher levels of APOE and larger, more lipidated extracellular particles at baseline and after NPC1 inhibition, whereas APOE4 astrocytes produce smaller particles that fail to mobilize lipid stores. Efficient lipid buffering by APOE3 astrocytes may help maintain neuronal membrane homeostasis under cholesterol stress, supporting cytoskeletal organization and normal APP anchoring in cholesterol-rich microdomains and γ-secretase cleavage patterns. In contrast, impaired lipid export by APOE4 astrocytes may exacerbate cholesterol accumulation, promoting aberrant APP processing, reduced Aβ40 production, and elevated Aβ42/40 ratios. These changes are predicted to ultimately compromise neuritic integrity. While not a primary objective of this study, we observed that cholesterol and lipid accumulation altered APP metabolism in the APOE3 and APOE4 neuron-astrocyte co-cultures, including increased APP-FL, elevated APP-CTFs, and a higher Aβ42/40 ratio. This is consistent with prior reports in NPC disease and in NPC1-deficient models where cholesterol buildup disrupted membrane microdomains and enhanced γ-secretase-dependent generation of longer Aβ species (5, 58, 59). We have previously shown that NPC1 inhibition in human cells increases levels of APP and BACE1 and drives APP association with neutral lipids, which is prevented by APOE2 and APOE3, but not APOE4. In the current study, APOE genotype did not abolish APP upregulation, but APOE3 astrocytes suppressed the accumulation of APP cleavage products, consistent with a buffering effect on membrane lipid composition and trafficking (5, 15, 57, 60). Together, these findings support a model in which cholesterol and lipid imbalances drive APP biochemical changes. Although much work has centered on investigations into the amyloid pathway, earlier studies clearly pointed to cholesterol and APOE regulation as central drivers (61, 62), yet this has remained understudied until recently. In the present experiments we also observed a selective reduction in Aβ40 upon NPC1 inhibition, with Aβ42 remaining stable. This is notable, as Aβ42 is classically viewed as the toxic species, yet accumulating evidence suggests that Aβ40 contributes to normal synaptic physiology. For example, elevations in endogenous Aβ, including Aβ40, enhance synaptogenesis and excitatory transmission in human neurons, whereas APP deletion or BACE1 inhibition suppressed synapse density and miniature EPSCs (63). Thus, disruptions in Aβ40 balance, rather than simple accumulation of Aβ42, may impair synaptic support even in the absence of aggregation. Proteomic analyses of amyloid plaque-associated regions in CRND8 mice, which overexpress mutant human APP, show that differentially expressed proteins include lipid-binding proteins and regulators of synaptic vesicles (64). Consistent with this, NPC1-related lipid accumulation may stabilize APP within cholesterol-rich microdomains; and because γ-secretase also processes other substrates such as NOTCH while APP metabolism is closely linked to trophic and cholinergic signaling (65), these changes could bias cleavage toward APP and contribute to the selective reduction in Aβ40 and altered Aβ42/40 balance we observe.

### Lipid dysregulation emerges as a central driver of neuronal vulnerability across multiple neurodegenerative diseases

Cholesterol and lipid imbalances are now emerging as a central theme and nexus of neuronal degeneration across multiple neurodegenerative diseases. In NPC1 models of NP disease, cholesterol and sphingolipids accumulate in lysosomes and the trans-Golgi network, disrupting vesicle trafficking and triggering APP misprocessing, tau abnormalities, and cognitive decline (5, 6, 58, 59, 66). Comparable disturbances occur in GBA1 carriers, where impaired lysosomal function and glycosphingolipid metabolism confer major risk for PD and LBD (2, 3). APOE4 carriers likewise exhibit deficits in lipid transport and particle formation that exacerbate neuronal stress (48, 67). Cholesterol imbalance has also been implicated in ALS, further broadening the spectrum of lipid-associated neurodegeneration (21). These convergent findings indicate that lipid accumulation and trafficking defects act upstream in disease pathogenesis rather than as secondary consequences. By capturing neuronal lipid buildup, neuritic abnormalities, and altered APP processing, our coculture system highlights mechanisms common to both monogenic disease (NPC) and complex disorders (AD, PD, LBD, ALS) (5, 15, 21, 68).

These observations underscore the need for disease models that extend beyond reductionist mouse genetics. Transgenic animals rarely capture the combined influences of aging, lipid metabolism, mitochondrial stress, and protein aggregation that dominate human disease. Human iPSC systems allow precise control over neuronal–glial interactions and can model the cumulative burden of disrupted lipid exchange and cellular imbalance. Such approaches may better delineate mechanisms underlying disease initiation and progression, challenging interpretations based solely on knock-in/knockout animal data that often miss the multifactorial drivers evident in patients.

### Future Directions: Lipid Composition, Astrocyte Function, and Therapeutic Potential

Our findings indicate that astrocytic lipid support is critical for maintaining neuronal structure, and that disruption of this process contributes directly to neuritic pathology. A key next step is to define the biochemical composition of the lipid accumulations observed under NPC1 inhibition. Lipidomic analysis will be necessary to determine whether cholesterol, esters, oxysterols, or other lipid species are the drivers of dysfunction.

Astrocyte heterogeneity also needs to be considered. Distinct astrocyte subtypes show different capacities for lipid handling and stress responses (69, 70). It remains unclear whether certain populations are more sensitive to APOE4-related defects, or whether subtype-specific reprogramming might restore protective function. Single-cell analyses of APOE4 astrocytes under conditions of lipid and cholesterol imbalance may help identify candidate pathways for therapeutic targeting.

These mechanistic insights open therapeutic avenues aimed at restoring APOE3-like function. Enhancing ABCA1 activity (54, 71), correcting APOE4 misfolding (72–74), or using mimetic peptides such as 4F (5) have all shown promise in improving lipidation and normalizing APP metabolism. The current findings position cholesterol and lipid handling as central to neurodegeneration and therapeutic intervention. Our coculture system provides a human platform to test these interventions in a genotype-specific manner. Such interventions could include enhanced APOE lipidation and APOE particle formation and delivery to preserve brain and cell structural integrity and prevent lipid-driven pathology.

Finally, emerging large human cross-correlated lipidomic to brain disease risk can provide personalized medicine based on APOE genotype and specific neuronal responses to lipid stress. Continued in-depth research on lipid use, mobilization and transfer in human cell biological systems is likely to yield significant benefits to patients suffering from neurodegenerative diseases.

## Acknowledgements

This research was supported by NIH/NIA RF1AG080636, DoD HT94252310930, DoD HT94252310931, the Harold and Ronna Cooper Family, the Parkinson Research Foundation, the Orchard Foundation Consolidated Anti-Aging Foundation. EDB is supported by the Harold and Ronna Cooper Post-Doctoral Fellowship for Parkinson’s Disease Research. We thank Dr. Li-Huie Tsai (Massachusetts Institute of Technology) for generously sharing APOE3 and APOE4 iPSC lines used in the study. We are also grateful to Dr. Oliver Cooper for his insightful discussions.

---

**Figure S1.**
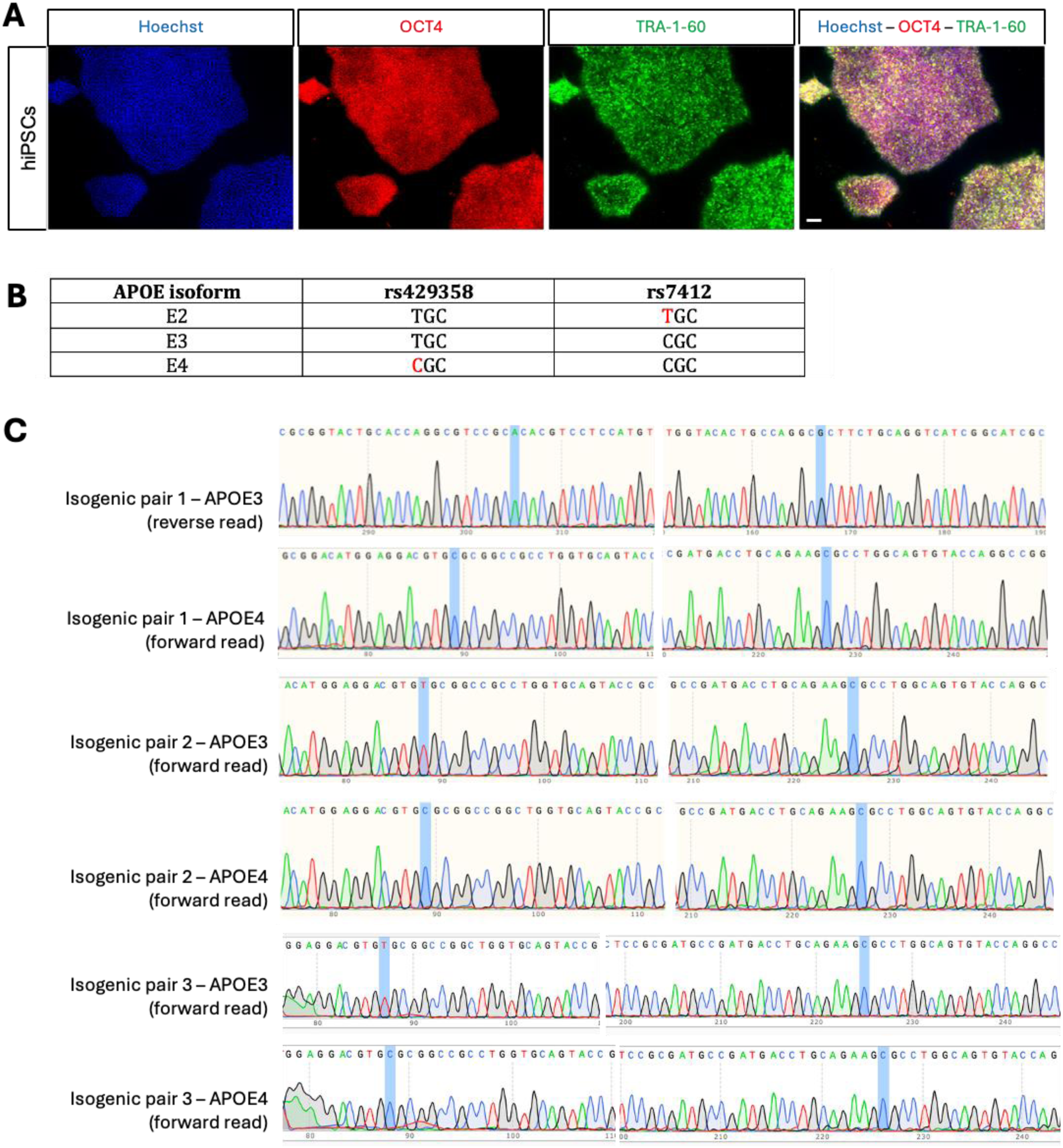
APOE genotyping of isogenic hiPSC pairs used in this study. (A) hiPSCs expressing pluripotency markers OCT4 (red) and TRA-1-60 (green). Nuclei were stained with Hoechst (blue). Scale bar: 100 μm. (B) The two single nucleotide polymorphisms (SNPs) that distinguish the 3 APOE isoforms are shown: rs429358 (T>C) and rs7412 (C>T). (C) Sanger sequencing confirms the presence of APOE3 or APOE4 SNPs in 3 isogenic hiPSC pairs.

**Figure S2.**
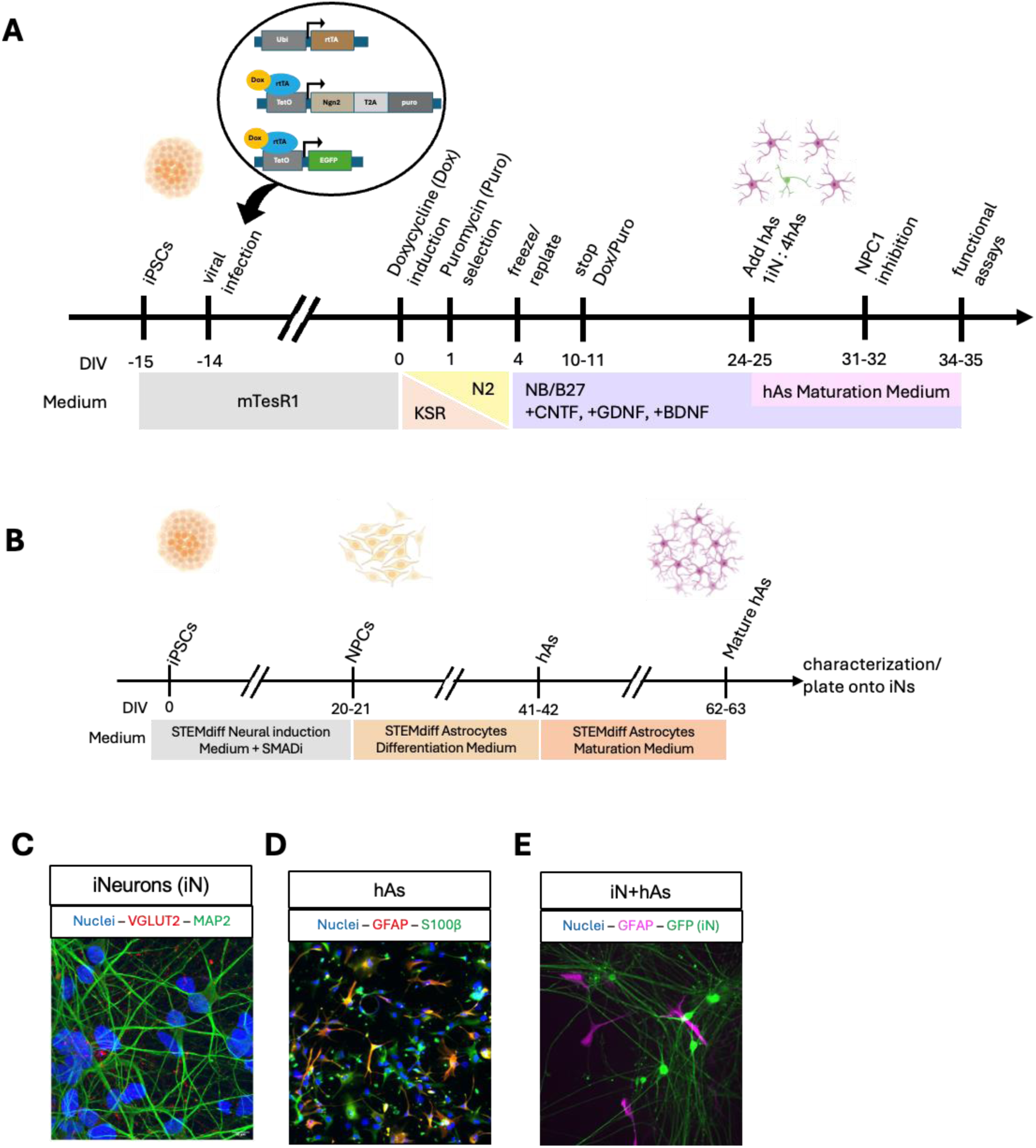
Generation of APOE3 and APOE4 iN and hAs from hiPSCs. (A) Schematic of neuronal differentiation. Excitatory glutamatergic iNeurons (iN) were generated from APOE3 or APOE4 hiPSCs by lentiviral transduction of NGN2 and GFP. NGN2 expression was induced with doxycycline, and transduced cells were selected with puromycin. Following 2 weeks of differentiation, iN were cocultured with astrocytes for 1 week and subsequently treated with the NPC1 inhibitor for 3 days to induce lipid stress. (B) Schematic of astrocyte differentiation. APOE3 or APOE4 hiPSCs were first differentiated into neural progenitor cells (NPCs) using STEMdiff™ Neural Induction Medium supplemented with SMAD inhibitors. NPCs were then cultured in STEMdiff™ Astrocyte Medium for 3 weeks, followed by an additional 3 weeks in STEMdiff™ Astrocyte Maturation Medium to obtain mature astrocytes (hAs). (C-E) Representative immunofluorescence images of cell type-specific markers. (C) Excitatory glutamatergic iN labeled with VGLUT2 (red) and the pan-neuronal marker MAP2 (green), (D) Mature hAs expressing astrocyte markers GFAP (red) and S100β (green), (E) iN+hAs cocultures, where GFP (green) marks iN, and GFAP (magenta) marks hAs.

